# Simultaneous Native Mass Spectrometry Analysis of Single and Double Mutants to Probe Lipid Binding to Membrane Proteins

**DOI:** 10.1101/2023.09.19.558516

**Authors:** Hiruni S. Jayasekera, Farhana Afrin Mohona, Megan Ewbank, Michael T. Marty

## Abstract

Lipids are critical modulators of membrane protein structure and function. However, it is challenging to investigate the thermodynamics of protein-lipid interactions because lipids can simultaneously bind membrane proteins at different sites with different specificities. Here, we developed a native mass spectrometry (MS) approach using single and double mutants to measure the relative energetic contributions of specific residues on Aquaporin Z (AqpZ) toward cardiolipin (CL) binding. We first mutated potential lipid-binding residues on AqpZ, and mixed mutant and wild-type proteins together with CL. By using native MS to simultaneously resolve lipid binding to the mutant and wild-type proteins in a single spectrum, we directly determined the relative affinities of CL binding, thereby revealing the relative Gibbs free energy change for lipid binding caused by the mutation. Comparing different mutants revealed that the W14 contributes to the tightest CL binding site, with R224 contributing to a lower affinity site. Using double mutant cycling, we investigated the synergy between W14 and R224 sites on CL binding. Overall, this novel native MS approach provides unique insights into lipid binding to specific sites on membrane proteins.

## Introduction

Lipids can modulate the structure, function, and stability of membrane proteins through different types of interactions.^[1–3]^ Indirectly, lipids can affect membrane proteins by changing the physical properties of the membrane, such as thickness or fluidity.^[4–7]^ Conversely, lipids can also directly interact with specific sites on the membrane protein to modulate the stability and activity of the protein.^[1,4–6,8,9]^ In between these two extremes, there is a sliding scale of specificities for lipids at different membrane protein binding sites, ranging from highly specific to totally nonspecific.^[10,11]^ The diversity of lipid interactions makes it challenging to quantify lipid binding affinity and selectivity at specific binding sites.

Native mass spectrometry (MS) has become a valuable tool for studying membrane protein-lipid interactions.^[12–15]^ By using non-denaturing ionization conditions, native MS preserves folded structures and noncovalent interactions for mass analysis.^[16–19]^ The ability of native MS to detect isolated membrane protein-lipid complexes has enabled measurement of lipid binding affinities and thermodynamics of protein-lipid interactions.^[15,20–23]^ However, the conventional native MS approach requires multiple titrations with varying lipid concentrations, which is slow, laborious, and prone to errors.

Unlike most other biophysical approaches, native MS can resolve small mass differences in a single spectrum, allowing lipid binding to be probed independently to any species that can be resolved in the mass spectrometer.^[17,24,25]^ Leveraging this power here, we developed a native MS experiment design to simultaneously examine the energetic contributions of selected amino acids to lipid binding by comparing wild-type and mutant proteins in the same spectrum. This single-mutant approach directly reveals differences in lipid binding affinity caused by the mutations, enabling ranking of lipid binding sites based on their thermodynamic trends.

Double mutant cycling was previously developed to determine the energetic coupling between amino acids in proteinligand^[26–28]^ and protein-protein interactions.^[29–34]^ In a double mutant cycle, two different amino acid residues are mutated separately and in combination to create individual mutants and their corresponding double mutant.^[29,31]^ The binding free energy is then measured for the wild type, each mutant, and the double mutant against each other. If the sum of the individual free energy changes for both single mutants relative to the wild type differs from the free energy change for the wild-type relative to the double mutant, the two residues are energetically coupled.^[17,31,35]^ Coupling between residues is a separate concept from coopertivity in binding. Cooperativity refers to binding of one ligand influencing the binding affinity of a second ligand, either positively or negatively. In contrast, coupling refers to how two residues bind a single ligand, either working together to promote binding or working in opposition. A system with a single binding site can have coupling but cannot have cooperativity. Systems with multiple binding sites can have both, but here we have focused exclusively on coupling.

Previous work by Horovitz and Sharon has used native MS to examine protein-protein interactions through double mutant cycle analysis.^[33]^ Because its high resolution can distinguish distinct masses of all the mutants, native MS provides a powerful advance of the method by enabling simultaneous detection of all interactions from a single spectrum.^[33]^ Here, we apply native MS to extend double mutant cycling to study the coupling between amino acid residues for membrane protein-lipid interactions.

We applied these advanced experiments to study the binding of cardiolipin (CL) (Figure S1) to aquaporin Z (AqpZ). AqpZ, a homo-tetramer, is responsible for the selective, passive transport of water across membranes.^[36–39]^ Previous work has shown that CL binds to AqpZ and affects its activity.^[40,41]^ A prior study by Schmidt and coworkers investigated a mutation that disrupted CL binding using native MS and molecular dynamic simulations, which proposed CL binding sites at the monomer-monomer interface of AqpZ.^[40–42]^ However, it remains challenging to identify and quantify the affinity of CL to specific binding sites and to measure the thermodynamics of CL binding to AqpZ.

Here, we measured the thermodynamic contributions of a range of amino acid residues on AqpZ-CL interactions using native MS on mixtures of AqpZ wild type, single mutants, or double mutants bound to CL. Our data revealed the relative change in the binding free energy caused by mutating specific residues on multiple CL-bound states. Moreover, double mutant cycle analysis revealed energetic coupling between two residues for CL binding.

### THEORY

We begin by considering the dissociation of a protein–lipid complex, *PL*, into free protein, *P*, and free lipid, *L*: *PL* ⇌ *P* + *L*. We can write a conventional expression for the dissociation constant, *K*_*D*_:

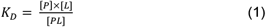

Using the conventional approach, if we assume that *P* and *PL* have similar detector responses, we can use native MS to measure the ratio of the signal intensities of *P* and *PL* from a single spectrum, but multiple spectra need to be collected to fit the unknown free lipid term, *L* .^[43,44]^ To compare the relative affinities of mutant and wild-type proteins using this approach, multiple titration series^[28]^ would have to be performed with each protein independently, increasing the time, labor, and potential for experimental errors.

Here, we developed an alternative approach to directly measure the relative binding affinities of two species from a single spectrum. First, both wild-type (*W*) and mutant (*M*) proteins are mixed with lipids in a single tube to form *WL* and *ML* complexes. Because all species are present in the same solution, the free lipid term, *L*, will be identical for both species. Thus, we can express the relative affinities of the wild-type (*K*_*D,w*_) and mutant (*K*_*D,M*_) protein as a ratio, *K*, that cancels out the free lipid term:

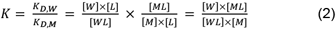

As noted above, we assume the native MS signal intensities are proportional to concentration for all terms in this equation. This ratio of dissociation constants for the *WL* and *ML* complexes is also the equilibrium constant for the exchange of a lipid between the mutant and wild type, *WL* + *M* ⇌ *W* + *ML*. Up to this point, we have focused on a single lipid binding the apo protein, but similar equations can be written for binding of a second lipid to a complex with one lipid bound, binding a third lipid to a complex with two, and so on.

Importantly, native MS can distinguish the wild-type and mutant proteins with different numbers of bound lipids in the same spectrum (Figure 2B and 1C), providing direct measurements of all terms in Equation 2 without the need for multiple titrations. This experiment design also has several inherent internal controls that limit experimental errors. Specifically, small errors in the lipid concentration are internally controlled because everything is combined in a single tube. Similarly, small errors in the protein concentrations do not factor into the equation because only the ratio of bound and unbound for each species is used.

**Figure 1.**
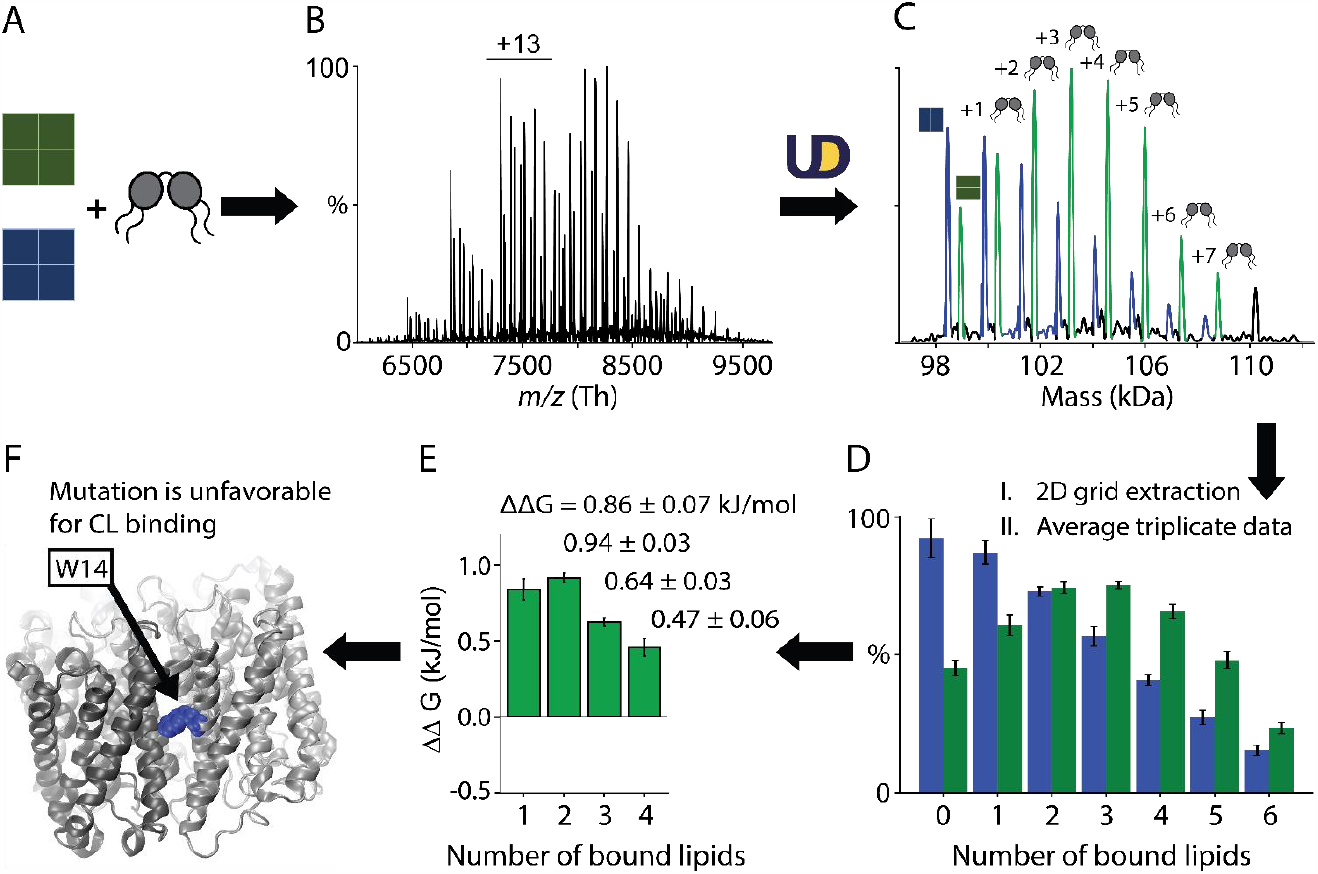
Schematic of single mutant analysis experiment and its data processing workflow. (A) Wild-type (WT) and W14A mutant proteins are mixed with CL at an approximate ratio of 1:1:100 of WT:W14A:CL. Wild type and W14A are annotated as tetrameric squares colored in *green* and *blue*, respectively, and CL is represented in a grey graphic. (B) Raw and (C) deconvolved mass spectra for a representative replicate at 25 °C for the wild type and mutant proteins with a series of lipids bound where deconvolved peaks are colored in *green* and *blue*, respectively. The number of lipids bound is annotated. Each peak area was extracted, and (D) the bar chart shows the average and standard deviation from the three replicate measurements. Calculated from Equation 3, (E) the differences in the Gibbs free energy change (ΔΔ*G*) plot for binding up to four lipids reveal that (F) the W14A mutation is unfavorable for CL binding. The W14 site on one monomer is annotated in *blue* while the entire protein complex is *grey*.

**Figure 2.**
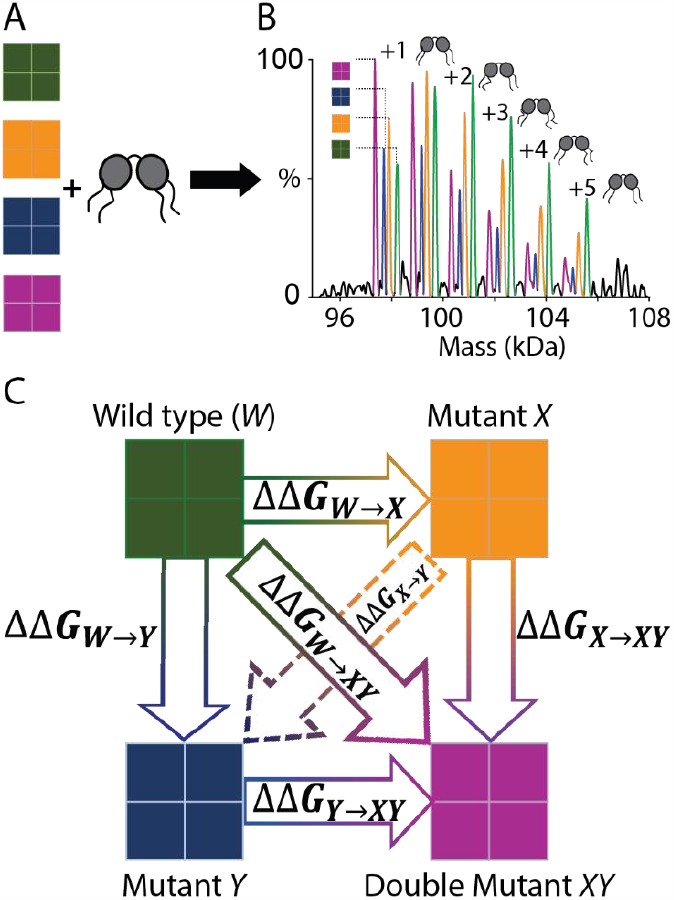
Schematic of the double mutant cycle experiment and its data processing workflow. (A) Wild-type and mutant proteins (*X, Y*, and *XY*) are mixed with CL at an approximate ratio of 1:1:1:1:200, where the proteins are represented in tetrameric squares colored in *green, yellow, blue*, and *purple*, respectively, and CL is represented in a *grey* graphic. (B) Deconvolved mass spectra independently resolve all four proteins, indicated in their respective colors. The numbers of bound lipids are annotated. (C) Extraction and pairwise comparison of the relative peak heights for each bound lipid enables the construction of the double mutant cycle.

From the equilibrium constant for the exchange of a lipid between the mutant and the wild-type forms, we can derive an expression for the difference in the Gibbs free energy change for lipid binding between wild type and mutant, ΔΔ*G* as:

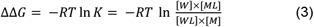

where *R* is the ideal gas constant, *T* is the temperature in Kelvin, and *K* is the equilibrium constant from Equation 2. The ΔΔ*G* indicates how favorable CL binding is to the mutant relative to the wild type. Positive values (Figure 1E) show the mutation is less favorable for CL binding to the mutant relative to the wild type, and negative values indicate the mutant is more favorable for CL binding. Therefore, the ΔΔ*G* values quantify how much each residue contributes to CL binding relative to the mutant form.

It is important to note that AqpZ is a homotetramer, and each mutation will thus affect four symmetric binding sites. Moreover, there are multiple sites where lipids can bind, which we have not tried to model. Thus, these ΔΔ*G* values do not measure a single lipid binding a single site. Rather, they represent the relative global binding affinities averaged across multiple sites and are most useful when considering the overall trends rather absolute energies. For simplicity, we have also not tried to examine lipid coopertivitiy but instead simply report results for binding of each lipid.

Combining sets of data for individual mutants with their double mutant to construct a double mutant cycle reveals cooperative interactions between residues during lipid binding. However, comparing data from single mutant analyses performed in separate tubes for each mutant with the wild type did not provide accurate energy differences, potentially because the protein mixing ratios and free lipid concentrations could vary between experiments. To overcome this limitation, we carried out a separate experiment with four proteins–wild type (*W*), mutant *X*, mutant *Y*, and the double mutant (*XY*)–in the same tube to construct the double mutant cycle (Figure 2).

For independent residues in this cycle that are not coupled, the ΔΔ*G* for a single mutation should be identical, regardless of whether the other mutation has been made: ΔΔ*G*_*W*→*X*_ − ΔΔ*G*_*Y*→*XY*_ = ΔΔ*G*_*W*→*Y*_ − ΔΔ*G*_*X*→*XY*_ = 0 (Figure 2C). A non-zero value indicates that there is an interaction or energetic coupling between the two mutated residues.^[45]^ Coupling can also be observed by comparing the free energy associated with the double mutant against the sum of independent free energies of individual mutants. In an uncoupled system, ΔΔ*G*_*W*→*XY*_ = ΔΔ*G*_*W*→*X*_ + ΔΔ*G*_*W*→*Y*_ (Figure 2C).^[35]^ In other words, if the double mutant is not equal to the sum of the parts, the two residues are coupled.

## Results and Discussion

### SINGLE MUTANT ANALYSIS

First, we investigated the thermodynamics of CL binding to different potential lipid binding sites on AqpZ by measuring simultaneous lipid binding to wild-type and mutant proteins using native MS. We began with the W14A mutation, which sits at the monomer interface (Figure 3) and was suggested to be involved in CL binding.^[39,40,42]^ After incubating the wild type and W14A mutant with CL, we performed native MS. In the spectra (Figure 1B, 1C, and S2), we observed a peak series for the wild-type and mutant proteins, with the mutant slightly lower in mass (Table S1). Within each peak series, distinct peaks were observed for up to 7 bound CL molecules, with each peak separated by the mass of CL.

**Figure 3.**
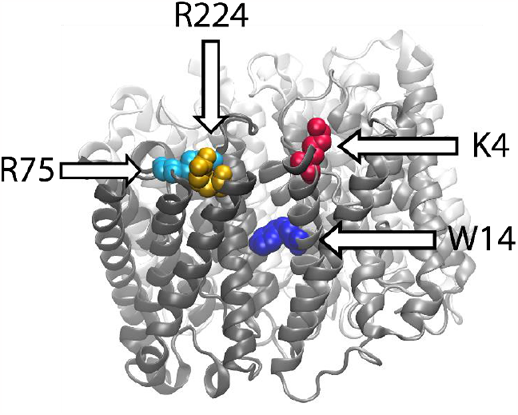
AqpZ with mutant sites W14, R224, R75, and K4 labeled, indicated with an arrow, and colored in *blue, yellow, cyan*, and *red*, respectively. Four chains of the protein are shown in *grey*. PDB code: 2abm.^[46]^

Globally, the peak series is clearly shifted towards lower lipid binding with the W14A mutant relative to the wild type (Figure 1C), which suggests a lower affinity of the mutant for CL binding. To quantify the relative affinity difference for each bound lipid, we calculated the equilibrium constant for lipid exchange between the mutant and wild type (Equation 2), and we used this value to calculate the change in Gibbs free energy for lipid binding caused by the mutation (Equation 3).

The positive ΔΔ*G* values for each of the first four bound lipids (Table S2 and Figure 4A) indicate that CL binding is less favorable to the W14A mutant than the wild type. In other words, the W14 residue contributes an average of about 0.86 kJ/mol to the global binding of the first CL, relative to an alanine in the same location. Interestingly, the effect was most pronounced when comparing 0 vs. 1 and 1 vs. 2 bound lipids. Comparing states with higher amounts of bound lipids, such as 2 vs. 3, we see positive ΔΔ*G* values but with decreasing magnitudes. When comparing even higher amounts of lipids bound, such as 4 vs. 5 and 5 vs. 6, the values continue to decrease, but confidence intervals increase at higher values due to the weaker signals (Figure S3).

**Figure 4.**
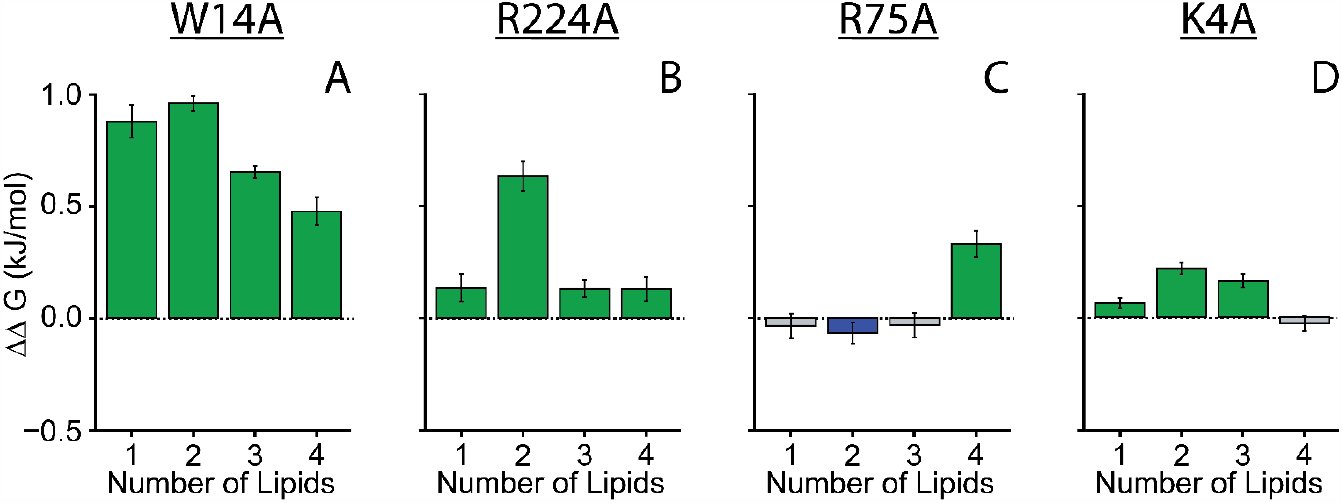
The difference in the Gibbs free energy change (ΔΔG plots) for CL binding up to four lipids at 25 °C for each mutant compared to the wild-type protein. Error bars indicate the 95% confidence intervals. *Green* color bars represent unfavorable mutations, where CL binding is more favorable to the wild type. Negative *blue* color bars depict more favorable CL binding to the mutant. *Grey* bars represent statistically insignificant interactions.

These data indicate that W14 participates either directly in a binding site for CL or plays an indirect role in the binding of CL to another site on AqpZ. Our interpretation for the observed decrease in ΔΔ*G* as more lipids bind is that the binding site involving W14 has the strongest affinity for CL, out of the residues studied here. If only one CL is bound (comparing 0 vs. 1), W14 makes a strong contribution to lipid binding at this higher affinity site. Because AqpZ is a homo-tetramer, there are 4 potential binding sites that involve W14. As additional lipids bind (1 vs. 2, 2 vs. 3, etc.), they can bind at the other open symmetric sites involving W14. However, they can also bind at different sites, due in part to their being fewer open binding sites involving W14 to access. Because these other binding sites are not probed by the W14A mutation, the W14A mutation has a lower contribution to the global free energy of lipid binding as more lipids are bound in more diverse sites.

After examining W14, we next explored surface exposed amino acids near the monomer interfaces of AqpZ. Because CL is anionic (Figure S1), we focused on cationic amino acids. We started with R224A mutant (Figure 3). We performed a single mutant analysis with wild-type and R224A mutant proteins mixed with CL. Like W14A, we observed distinct peak series for the wild-type and R224A proteins with up to 7 lipids bound (Figure S2), and we calculated positive ΔΔ*G* values (Table S2 and Figure 4B), which indicates that CL binding was also less favorable to the R224A mutant than the wild type.

However, the ΔΔ*G* values for R224A were lower than the values for W14A, revealing a higher overall CL binding affinity from the sites involving W14 over sites involving R224. Interestingly, for wild type and R224A, the highest ΔΔ*G* value was for the 2^nd^ lipid binding (1 vs. 2 bound lipids). Building on our interpretation of the W14A data discussed above, we interpret this pattern to indicate that the site involving R224 is the second highest affinity site for CL binding on AqpZ. When the first CL binds (0 vs. 1), it primarily binds the site involving W14. When the second lipid binds (1 vs. 2), it can bind at either one of the three open sites involving W14 or one of the four open sites involving R224. Although the W14 site is preferred, as evidenced by the statistically higher ΔΔ*G* between 1 vs. 2 CL bound for W14A relative to R224A, there is still a significant population of lipids that bind at the R224 site after the first W14 site is occupied (compare lipid 2 in Figure 4A and B).

We then explored the R75 residue, which is close to R224 on the surface of AqpZ near the monomer interface (Figure 3). Single mutant analysis reveals that this mutant binds CL with slightly higher affinity than the wild type, as evidenced by negative ΔΔ*G* value (Table S2 and Figure 4C). However, these data indicate that R75 site does not contribute as much to CL binding. We only discovered a meaningful ΔΔ*G* when comparing 3 vs. 4 bound lipids. This observation suggests that the R75 residue is not important until at least 3 lipids are already bound to the complex, likely at the higher affinity sites involving W14 and R224.

Finally, we tested the K4 residue on the opposite side of the monomer interface from R224 (Figure 3). We performed a single mutant analysis between the wild-type protein and the K4A mutant in the same manner. K4A exhibited only minor effects on CL binding, with statistically positive but very small ΔΔ*G* values. Thus, the K4 residue does not contribute much to CL binding.

R224 and R75 are close to each other, so we wanted to compare the wild-type protein and the double mutant of R224A and R75A. Like the R224A mutant, the double mutant also exhibited lower CL binding at the 2^nd^ bound lipid, with the highest ΔΔ*G* value for 1 vs. 2 bound lipids (Table S2 and Figure S4). These results more closely resembled the R224A mutant, confirming that the R224 residue has a stronger contribution to CL binding than R75.

Overall, our single mutant analyses indicate that W14 contributes to the highest affinity lipid binding site, with R224 contributing to a second high affinity binding site. R75 only contributes to CL binding after several lipids are already bound, indicating a weak affinity, and K4 does not contribute much overall. Together, these data reveal multiple potential binding sites for CL around AqpZ and allow us to rank the binding sites based on the trends in their relative affinities.

## DOUBLE MUTANT CYCLE ANALYSIS

Based on our single mutant analysis, we propose that the W14 and R224 sites contribute to the highest and second highest affinity binding sites on AqpZ, respectively. To understand whether the binding of CL to one site (W14) influences the binding of CL to the other site (R224), we used double mutant cycle analysis, which allows us to explore the energetic coupling between residues.^[17,29,31–33,35]^

For the double mutant cycle experiment, we mixed all four proteins (wild-type, W14A, R224A, and R224A_W14A) in the same tube (Figure 2A). We observed resolved peaks with up to 5–6 bound lipids in all four proteins (Figure 2B–C, and Figure S5). Comparing the ΔΔ*G* plots obtained for wild-type with single mutant analysis (Figure 4A–B, and Table S2) and the double mutant cycle experiment (Figure S5 D–E and Table S3), we observed consistent patterns for the CL binding. However, there are slight differences in the values, potentially due to differences in free lipid concentrations and protein mixing ratios between experiments as well as influences from the other proteins.

We constructed the double mutant cycles for binding of each of the first four lipids by pairwise comparison of each species (Figure S6). However, we focused on the first two bound lipids (0 vs. 1 and 1 vs. 2) because those were most affected by these mutations (Figure 5). In both cases, the W14A mutation had a more pronounced effect on CL binding than the R224A mutation, with the left side greater than the top side of the box. We can directly quantify the difference in effects of the two mutants by comparing the upward diagonal between W14A and R224A. Here, we found that the W14A was 0.35±0.05 kJ/mol more disruptive than R224A for the first CL binding and 0.42±0.08 more disruptive for the second CL (Figure 5).

**Figure 5.**
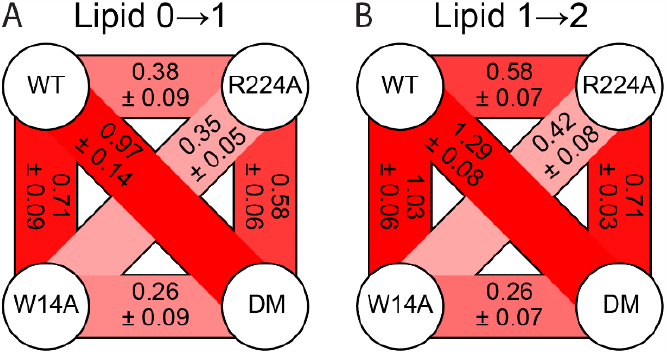
Double mutant cycle for the (A) first and (B) second CL bindings constructed with AqpZ WT, R224A, W14A, and R224A_W14A (DM) at 25 °C. The mean ΔΔ*G* in kJ/mol with ± 95% confidence intervals are stated. Positive ΔΔ*G* values are depicted in darker *red* for higher values. Data from lipids 2 to 3 and 3 to 4 can be found in Figure S6 C–D, respectively.

To investigate coupling between the W14 and R224 residues, we compared the ΔΔ*G* associated with the double mutant and the individual mutants. If the selected mutants are energetically independent, ΔΔ*G* for the double mutant would be the sum of the ΔΔ*G* values for the individual mutants: ΔΔ*G*_*W*→*XY*_ = ΔΔ*G*_*W*→*X*_ + ΔΔ*G*_*W*→*Y*_ (Figure 2C). In simpler terms, the diagonal of the box in Figure 5A would be equal to the sum of the top and left sides if the residues were independent. For the first bound lipid (0 vs. 1), we discovered that the ΔΔ*G* between the wild type and double mutant (0.97 ± 0.14 kJ/mol) was statistically equal to the sum of the individual mutants (1.09 ± 0.13 kJ/mol), indicating independent CL binding at each site.

Similarly, independent free energies should show the same ΔΔ*G* for the same mutation, regardless of whether the other residue has been mutated. In other words, opposite sides of the box in Figure 5A should be equal if the residues are independent. For the first bound lipid (0 vs. 1), we found that the differences in ΔΔ*G* values between the top arm vs. bottom for the R224A mutation (0.12 ± 0.12 kJ/mol) and the right arm vs. left for the W14A mutation (0.13 ± 0.11 kJ/mol) were small, indicating again that these sites do not appear to interact for the first lipid binding at high concentrations.

However, for the second lipid binding on AqpZ (Figure 5B), the two residues were not independent but instead couple. The diagonal comparing the double mutant to the wild type (1.29 ± 0.08 kJ/mol) was not equal to the sum of the individual mutants (1.61 ± 0.09 kJ/mol). Instead, the double mutant was lower than the sum of the two single mutants, indicating a coupling energy, *CE* = ΔΔ*G*_*W*→*XY*_ − (ΔΔ*G*_*W*→*X*_ + ΔΔ*G*_*W*→*Y*_), of −0.32 ± 0.12 kJ/mol between the W14 and R224 residues for CL binding, relative to alanines at the same locations. Coupling was also evident when considering the two opposing sides of the box (top and bottom, or right and left), which showed a difference of around 0.32 kJ/mol between each pair of arms.

To determine whether cooperativity is positive or negative, we considered the change in CL binding from each mutation. The impact of the W14A mutation was –0.32 ± 0.07 kJ/mol less impactful when the R224A mutation was already present (right arm < left arm). This difference shows that the W14A mutation was less detrimental to CL binding if the R224A mutation was already made. Similarly, the R224A mutation was –0.32 ± 0.10 kJ/mol less impactful when the W14A mutant was already made (bottom arm < top arm). Because the effect of one mutation became less pronounced when the other mutation was present, the presence of both residues is more favorable to CL binding than expected from each single residue, a positive synergy in binding CL. Therefore, R224 and W14 work together to promote CL binding to AqpZ.

Finally, we investigated the concentration dependence of these changes by varying the membrane protein:lipid ratio. At lower CL concentrations, there were not enough lipids to detect three or four bound lipids (Figure S7 and Table S3). However, for the first and second bound lipid, the W14A mutation became more dominant than R224A (Figures S8 and S9). These data confirm that R224 contributes to a lower affinity binding site that is less occupied at lower concentrations than the higher affinity site involving W14.

Interestingly, the coupling between W14 and R224 was stronger for the first lipid at lower concentrations, despite the lower independent contributions from R224 (Figures S8, S9, and Table S3). Our interpretation is that the R224 residue can contribute weakly to promote CL binding at the high affinity site. These contributions are entirely dependent on the presence of W14, as evidenced by the negligible differences between the double mutant and the W14A mutant. In other words, when W14 is mutated, R224 no longer contributes. At higher concentrations, this coupling is less prominant because other lower affinity binding sites can be populated, including a lower affinity site with independent R224 contributions. At lower concentrations, only the high affinity binding site is probed, so coupling may be more easily detected.

From a structural perspective, these data indicate that there is a higher affinity CL binding site involving W14. R224 contributes weakly to this higher affinity binding site in a manner that is entirely dependent on the presence of W14. However, R224 can also contribute independently to a lower affinity binding site that becomes more occupied at higher concentrations. It is not clear if coupling energies observed for the second bound lipid also indicate that binding of an initial CL at a the higher affinity W14 site promotes binding of second CL at the lower affinity site involving independent R224 (inter-site coupling). In any case, our results reveal that membrane protein residues can work together to facilitate lipid binding to membrane proteins.

## Conclusion

To study the contributions of specific residues to the thermodynamics of membrane protein-lipid interactions, we used native MS to simultaneously measure lipid binding to a pair or quartet of membrane protein mutants. We simultaneously determined the relative binding affinities of CL to both wild type and single mutants, which enabled us to identify W14 as contributing to the highest affinity binding site and R224 as contributing to the second highest affinity binding site. In contrast, R75 and K4 sites do not play a major role in binding to CL.

To investigate coupling interactions between W14 and R224 residues through double mutant cycling, we simultaneously mixed all four proteins: wild-type, two single mutants, and their corresponding double mutant. Interestingly, we observed a positive cooperative interaction between the two residues for CL binding.

In contrast to other techniques, native MS with mutant analyses offers a simple method for comparing thermodynamics of intermolecular interactions from a single spectrum.^[33]^ Future work will explore thermodynamic parameters such as enthalpy and entropy through Van ‘t Hoff analysis, which unfortunately was not achievable in this study.^[15]^ It may also be possible to build on these experiments to extract more complex information on the affinities of individual microscopic binding sites and on lipid binding cooperativity. Although our simple binding model does not capture all the complexity of lipid binding to membrane protein surfaces, this method provides valuable insights into understanding membrane protein-lipid interactions. Specifically, the trends in binding help rank binding sites by their relative affinities and binding patterns. The measurements also reveal the presence and direction of coupling.

## Supporting Information

The authors have cited additional references within the Supporting Information.^[47–52]^

## Acknowledgements

The authors thank Maria Reinhardt-Szyba, Kyle Fort, and Alexander Makarov at Thermo Fisher Scientific for support on the Q-Exactive HF UHMR instrument. We also thank Professors Amnon Horovitz, Michal Sharon, and attendees at the Advancing Mass Spectrometry Conference 2023 for their constructive remarks and insights. This work was funded by the National Institutes of Health (NIH) under grant numbers R35 GM128624 and RM1 GM145416 to M.T.M.

## Data availability statement

The data from this study are available in MassIVE: https://doi.org/doi:10.25345/C53R0Q44F.

## Entry for the Table of Contents

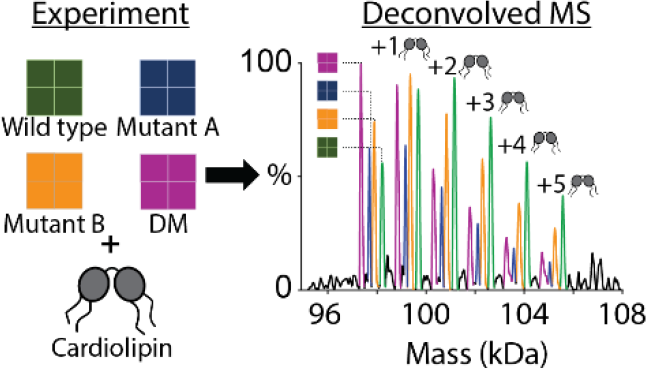

A single native mass spectrum with double mutant cycling reveals the energetic contributions of specific residues on lipids binding to membrane proteins and coupling between residues to promote lipid binding, uncovering new insights into the structure and thermodynamics of lipid binding to membrane protein complexes.

Institute and/or researcher Twitter usernames: @michaeltmarty @Biochem_Chem

