## Supporting Information for "Simultaneous Native Mass Spectrometry Analysis of Single and Double Mutants to Probe Lipid Binding to Membrane Proteins"

### Experimental Procedures

**Mutant Selection and Mutagenesis**

A previous molecular dynamics simulation study on Aquaporin Z (AqpZ) by Schmidt *et al.* found that CL tends to associate at the interface between monomers, and the occupancy of CL around AqpZ was reduced when W14 was mutated to alanine.[1] Therefore, we selected W14A as the first mutant. To further identify the specific CL binding sites, we chose positively charged arginine and lysine residues near the monomer-monomer interface, specifically R75, R224, and K4. Because CL is anionic (Figure S1), we predicted that it might interact with cationic residues.

Selected residues were mutated to alanine through site-directed mutagenesis to create single mutants, W14A, R224A, R75A, K4A, and double mutants, R224A_R75A and R224A_W14A (see Figure 3). Primers were ordered from Integrated DNA Technologies, Inc (San Diego, California, USA). Mutagenesis was performed by polymerase chain reaction (PCR) using Phusion® High-Fidelity PCR Master Mix with GC Buffer (New England Biolabs) using the AqpZ wild-type plasmid DNA as the template.[2–4] The DNA template was digested with DpnI (New England Biolabs). The mutant DNA was transformed into *E. coli* Stellar competent cells (Sigma Aldrich), and plasmids were isolated using a NucleoSpin® Plasmid kit (Macherey-Nagel) for confirmation by sequencing (Eton Bioscience, Inc).

**AqpZ Membrane Protein Expression and Purification**

Upon sequence confirmation, AqpZ wild-type and mutant plasmids were transformed into *E. coli* OverExpress C43 (DE3) cells (Sigma Aldrich). All forms of AqpZ-TEV-GFP-HIS were purified using immobilized metal affinity chromatography (IMAC) and size exclusion chromatography (SEC), as previously described.[4–6] Briefly, cells were cultured in flasks with 100 mL culture of Luria-Bertani (LB) media with 0.1 mg/mL Ampicillin and grown overnight at 37 °C while shaking at 200 rpm. The overnight culture was used to inoculate 6 flasks of 1 L Terrific Broth (TB) media (7 mL per 1 L of TB in a 2 L Erlenmeyer flask) with 0.1 mg/mL Ampicillin and was incubated until the optical density ($OD_{600}$) reached 0.6–0.8. Then, isopropyl β-D-1-thiogalactopyranoside (IPTG) was added to 1 mM to induce protein expression. After adding IPTG, the cells were incubated overnight at 24 °C. Cells were then harvested by centrifugation at 5,000 rpm for 10 min at 4 °C. Cells were lysed in 20 mM Tris, 0.1 M NaCl, pH 7.4 with protease inhibitor using an LM20 microfluidizer (Microfluidics International Corporation) at 20,000 psi. The cell lysate was clarified by centrifugation at 20,000 ×g for 25 minutes at 4 °C, and membranes were harvested at 100,000 ×g for 2 hours and 10 minutes at 4 °C. The membranes were resuspended in loading buffer (40 mM Tris, 0.3 M NaCl, 20 mM imidazole, 10% glycerol, 0.025% DDM, 5 mM β-mercaptoethanol (BME), pH 7.4 at room temperature (RT)) and homogenized.

AqpZ was then extracted from the purified membranes using 1% *n*-octyl-β-D-glucoside (OG) and *n*-dodecyl-β-D-maltoside (DDM) and incubated overnight in a tube rotator at 4 °C. Extracted membranes were clarified at 20,000 ×g for 25 minutes at 4 °C. The supernatant was filtered using 0.45 µM PES filters. AqpZ was purified using immobilized metal affinity chromatography (IMAC) using HisTrap HP (Cytiva) followed by concentration to 5 mL with a 100 kDa molecular weight cutoff (MWCO) concentrator (Sartorius Vivaspin™) and size exclusion chromatography (SEC) using a Superdex 200 16/600 (GE Healthcare) in the loading buffer. The purified AqpZ was incubated overnight with tobacco etch virus (TEV) protease at 4 °C to cleave the GFP-HIS tag. Cleaved protein was then purified by IMAC followed by the buffer exchange into 200 mM ammonium acetate buffer with 0.5% tetraethylene glycol monooctyl ether (C8E4) using SEC with a Superose 6 10/300 Increase GL (Cytiva). Peak fractions were concentrated to 1–10 µM with 50 kDa MWCO. Samples were flash frozen at -80 °C until the analysis.

**Native Mass Spectrometry (MS) Sample Preparation and Data Collection**

1',3'-bis[1-palmitoyl-2-oleoyl-*sn*-glycero-3-phospho]-glycerol (sodium salt) (16:0-18:1 CL) was purchased from Avanti Polar Lipids, dissolved in chloroform, quantified using phosphate analysis, and dried under $N_2$ gas and vacuum overnight.[1,2] A stock concentration of 1.5 mM CL was prepared in 200 mM ammonium acetate buffer with 0.5% C8E4, as previously described.[1] We performed another phosphate analysis on the stock CL prepared to confirm the exact concentration.

For the single mutant experiment, pair-wise combinations of wild type with single mutants and double mutants were mixed, and CL was added to each before incubation at 4 °C for 15–30 minutes. Membrane proteins were mixed at a roughly 1:1 molar ratio, but this ratio was slightly adjusted for each pair to achieve approximately equal MS signals for the two proteins. We used a ratio of approximately 1:1:100 protein A: protein B: CL, but the CL ratio was slightly adjusted between a range of 100–120 to achieve a spectral density of 6–7 bound lipids, which allowed us to measure multiple lipid-bound states.

For double mutant cycle analysis of two high-affinity amino acid residues, we mixed AqpZ wild type, R224A, W14A, and their corresponding double mutant (DM) at an approximately 1:1:1:1 ratio. This ratio was slightly adjusted to achieve approximately equal MS signals for all proteins. Then, we added CL prior to the incubation at 4 °C for 15–30 min to achieve an approximate ratio of 1:1:1:1:200 WT:R224A:W14A:DM:CL to obtain 5–6 bound lipids. To observe the concentration dependence, we also performed double mutant cycle experiment at lower CL ratios of 1:1:1:1:100, 1:1:1:1:50, and 1:1:1:1:25.

Native MS of the samples was performed using a Q-Exactive HF Orbitrap mass spectrometer (Thermo Fisher Scientific, Bremen) equipped with Ultra-High Mass Range modifications, as previously described.[4,5,7] The nano-electrospray source was equipped with a variable temperature (VT) source, as previously described.[8] Borosilicate needles were pulled with a P-1000 micropipette puller (Sutter Instrument, Novato, CA), and 4–6 µL of the sample was loaded into the needle for the MS analysis.[5] Upon placing the needle inside the VT source, the sample was allowed to equilibrate to the set temperature for 2 minutes. MS was performed in positive ion mode, and key MS settings included a mass range of 2000–25000 *m/z* for single mutant analysis and 4000–15000 *m/z* for double mutant cycle analysis, resolution at 15,000, 1.10–1.15 kV capillary voltage, 200 °C capillary temperature, 50–60 V source fragmentation, 80–95 V collision voltage, and trapping gas pressure setting of 7. Once the sample was equilibrated, mass spectra were acquired at the set temperature for 1.5 minutes. Similarly, spectra were acquired at temperatures ranging from 15–35 °C at 5 °C intervals.

To verify that AqpZ subunits with different mutants do not mix to form hybrid complexes (such as a tetramer complex with two mutant and two wild-type monomers), we mixed the wild-type and double mutant proteins without lipid and checked the mass spectrum over time. If subunit exchange were occuring, we would expect to see heterogeneous species with unique masses appearing in between the two homogeneous starting peaks. Instead, even after an overnight incubation at 4 °C, there was no evidence of subunit exchange (Figure S10). Nonetheless, it is advisable not to prolong incubation times for longer than necessary, as this may result in protein degradation and/or subunit exchange.

**Native MS Data Analysis**

MS data analysis was performed using UniDec[9] and custom Python scripts. Example scripts for data analysis can be found at: https://github.com/michaelmarty/UniDec/tree/master/PublicScripts/MutantCycleAnalysis.

Step 1: Mass deconvolution: Raw mass spectra were first deconvolved in UniDec. Main deconvolution parameters included a mass range of 96–112 kDa, and a charge range of 1–25, and the masses were sampled every 1 Da. The peak FWHM was set to 5 Th.

Step 2: Peak Area Extraction: After deconvolution, we used the 2D Grid Extraction tool in UniDec to extract the peak area for all wild-type and mutant peaks with lipids bound. With this tool, we specified the first protein mass (Mass 0), lipid mass (Mass 1), and the mass difference between the first and second protein masses (Mass 2), which corresponds to the amino acid change (see Table S1). We also set the minimum and maximum number of lipids (0–7) and mutations (0–1). For the extraction, we measured the peak areas above a 10% relative intensity threshold with a mass window of ±90 or 100 Da. Extracted peak areas were exported to a text file.

Step 3: $\Delta\Delta G$ Calculations: Using the intensities extracted from Step 2 for the mutant and wild-type forms, with and without lipids bound, the dissociation constants ($K$) were calculated using Equation 2 for each bound lipid from 1–6 at each temperature. Next, using the calculated $K$ values, $\Delta\Delta G$ values were calculated using Equation 3 for each temperature, each lipid bound state, and each pair of mutants. For higher accuracy, we performed Van 't Hoff analysis by plotting $\ln(K)$ against $1/T$. Linear regression of the curve was used to extract enthalpy and entropy values. Unfortunately, the errors on enthalpy and entropy were large and were thus not used further. Results were qualitatively the same for each temperature even at 37 °C and the average $\Delta\Delta G$ was determined by solving the linear regression at 25 °C, and their corresponding 95% confidence intervals are reported at this temperature (see Table S2 and Figure S3 for single mutant analysis and Table S3 and Figure S6 for double mutant cycle analysis).

### Supporting Tables

**Table S1.** Parameters used in extracting the deconvolved mass peaks using 2D grid extraction tool in UniDec. Mass 0 corresponds to the mass of the membrane protein with the lower mass, mass 1 is the mass of CL, and mass 2 corresponds to the mass difference between membrane protein A and B.

| Membrane Protein A | Membrane Protein B | Mass 0 (Da) | Mass 1 (Da) | Mass 2 (Da) |
|---|---|---|---|---|
| WT | W14A | 98,405 | 1,410 | 485 |
| WT | R224A or R75A | 98,590 | 1,410 | 300 |
| WT | K4A | 98,700 | 1,410 | 225 |
| WT | R224A_R75A | 98,270 | 1,410 | 655 |
| WT | R224A_W14A | 98,070 | 1,410 | 820 |
| W14A | R224A_W14A | 98,070 | 1,410 | 335 |
| R224A | R224A_W14A | 98,070 | 1,410 | 520 |
| R224A | W14A | 98,405 | 1,410 | 185 |

**Table S2.** Single Mutant Analysis. The mean difference in the Gibbs free energy change (ΔΔG (kJ/mol)) and the 95% confidence interval for CL binding between each pairwise membrane protein combination for up to four lipids at 25 °C.

| Membrane Protein A | Membrane Protein B | Lipid Number | Mean ΔΔG (kJ/mol) | Confidence Interval |
|---|---|---|---|---|
| WT | W14A | 1 | 0.86 | ±0.07 |
| WT | W14A | 2 | 0.94 | ±0.03 |
| WT | W14A | 3 | 0.64 | ±0.03 |
| WT | W14A | 4 | 0.47 | ±0.06 |
| WT | R224A | 1 | 0.13 | ±0.06 |
| WT | R224A | 2 | 0.62 | ±0.07 |
| WT | R224A | 3 | 0.13 | ±0.04 |
| WT | R224A | 4 | 0.13 | ±0.05 |
| WT | R75A | 1 | -0.03 | ±0.05 |
| WT | R75A | 2 | -0.07 | ±0.05 |
| WT | R75A | 3 | -0.03 | ±0.06 |
| WT | R75A | 4 | 0.33 | ±0.06 |
| WT | K4A | 1 | 0.06 | ±0.02 |
| WT | K4A | 2 | 0.22 | ±0.03 |
| WT | K4A | 3 | 0.16 | ±0.03 |
| WT | K4A | 4 | -0.03 | ±0.03 |
| WT | R224A_R75A | 1 | 0.00 | ±0.17 |
| WT | R224A_R75A | 2 | 0.70 | ±0.06 |
| WT | R224A_R75A | 3 | 0.15 | ±0.07 |
| WT | R224A_R75A | 4 | 0.05 | ±0.03 |

**Table S3.** Double mutant cycle analysis of AqpZ wild type, R224A, W14A, and their corresponding double mutant (DM) with CL at a 1:1:1:1:*n* molar ratio, where *n* is 25, 50, 100, and 200 of CL. The mean difference in the Gibbs free energy change (ΔΔG (kJ/mol)) and the 95% confidence interval for CL binding between pair-wise combinations of Mutant A and Mutant B, and the coupling energy between R224A and W14A mutants with the 95% confidence interval for up to four lipids at 25 °C at different CL ratios. Dash marks indicate values where the signal was too low to be accurately measured.

| Mutant A | Mutant B | Lipid Number | *n* = 200 | | *n* = 100 | | *n* = 50 | | *n* = 25 | |
|---|---|---|---|---|---|---|---|---|---|---|
| | | | Mean ΔΔG | Conf. Interval | Mean ΔΔG | Conf. Interval | Mean ΔΔG | Conf. Interval | Mean ΔΔG | Conf. Interval |
| WT | W14A | 1 | 0.71 | ±0.09 | 1.00 | ±0.09 | 1.11 | ±0.09 | 1.45 | ±0.15 |
| WT | W14A | 2 | 1.03 | ±0.06 | 0.92 | ±0.09 | 1.06 | ±0.09 | - | - |
| WT | W14A | 3 | 0.23 | ±0.08 | 0.46 | ±0.12 | - | - | - | - |
| WT | W14A | 4 | 0.08 | ±0.11 | - | - | - | - | - | - |
| WT | R224A | 1 | 0.38 | ±0.09 | 0.52 | ±0.05 | 0.37 | ±0.09 | 0.35 | ±0.06 |
| WT | R224A | 2 | 0.58 | ±0.07 | 0.38 | ±0.05 | 0.37 | ±0.07 | - | - |
| WT | R224A | 3 | 0.12 | ±0.05 | 0.20 | ±0.08 | - | - | - | - |
| WT | R224A | 4 | 0.07 | ±0.05 | - | - | - | - | - | - |
| WT | R224A_W14A | 1 | 0.97 | ±0.14 | 1.17 | ±0.08 | 1.11 | ±0.13 | 1.35 | ±0.08 |
| WT | R224A_W14A | 2 | 1.29 | ±0.08 | 0.94 | ±0.08 | 1.02 | ±0.14 | - | - |
| WT | R224A_W14A | 3 | 0.48 | ±0.11 | 0.65 | ±0.09 | - | - | - | - |
| WT | R224A_W14A | 4 | 0.31 | ±0.10 | - | - | - | - | - | - |
| W14A | R224A_W14A | 1 | 0.26 | ±0.09 | 0.21 | ±0.06 | 0.00 | ±0.11 | -0.10 | ±0.1 |
| W14A | R224A_W14A | 2 | 0.26 | ±0.07 | 0.02 | ±0.11 | -0.04 | ±0.12 | - | - |
| W14A | R224A_W14A | 3 | 0.23 | ±0.10 | 0.17 | ±0.12 | - | - | - | - |
| W14A | R224A_W14A | 4 | 0.24 | ±0.15 | - | - | - | - | - | - |
| R224A | R224A_W14A | 1 | 0.59 | ±0.06 | 0.65 | ±0.07 | 0.74 | ±0.11 | 1.00 | ±0.05 |
| R224A | R224A_W14A | 2 | 0.71 | ±0.03 | 0.56 | ±0.05 | 0.65 | ±0.10 | - | - |
| R224A | R224A_W14A | 3 | 0.36 | ±0.07 | 0.45 | ±0.09 | - | - | - | - |
| R224A | R224A_W14A | 4 | 0.23 | ±0.08 | - | - | - | - | - | - |
| R224A | W14A | 1 | 0.35 | ±0.05 | 0.34 | ±0.09 | 0.78 | ±0.07 | 1.03 | ±0.12 |
| R224A | W14A | 2 | 0.42 | ±0.08 | 0.51 | ±0.10 | 0.78 | ±0.08 | - | - |
| R224A | W14A | 3 | 0.11 | ±0.08 | 0.31 | ±0.16 | - | - | - | - |
| R224A | W14A | 4 | -0.02 | ±0.12 | - | - | - | - | - | - |
| Coupling Energy | | 1 | 0.10 | ±0.18 | 0.35 | ±0.13 | 0.37 | ±0.18 | 0.45 | ±0.18 |
| Coupling Energy | | 2 | 0.32 | ±0.11 | 0.36 | ±0.13 | 0.41 | ±0.16 | - | - |
| Coupling Energy | | 3 | 0.13 | ±0.08 | 0.01 | ±0.17 | - | - | - | - |
| Coupling Energy | | 4 | 0.15 | ±0.12 | - | - | - | - | - | - |

### Supporting Figures

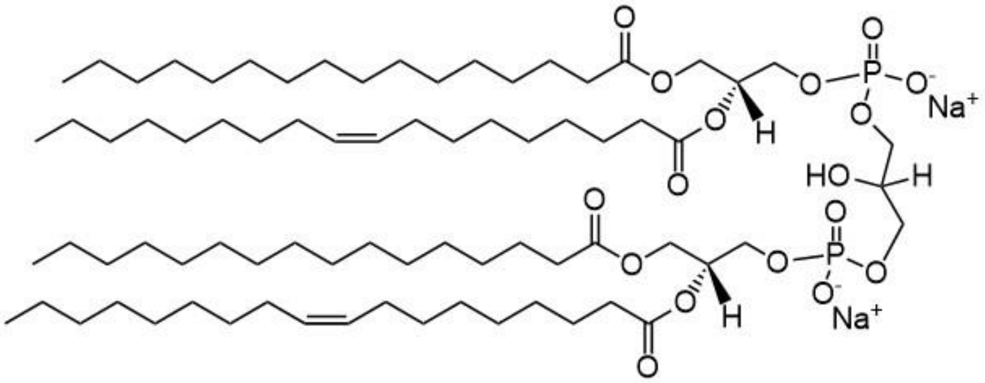

**Figure S1.** Structure of 1',3'-bis[1-palmitoyl-2-oleoyl-*sn*-glycero-3-phospho]-glycerol (sodium salt) (16:0-18:1 CL).

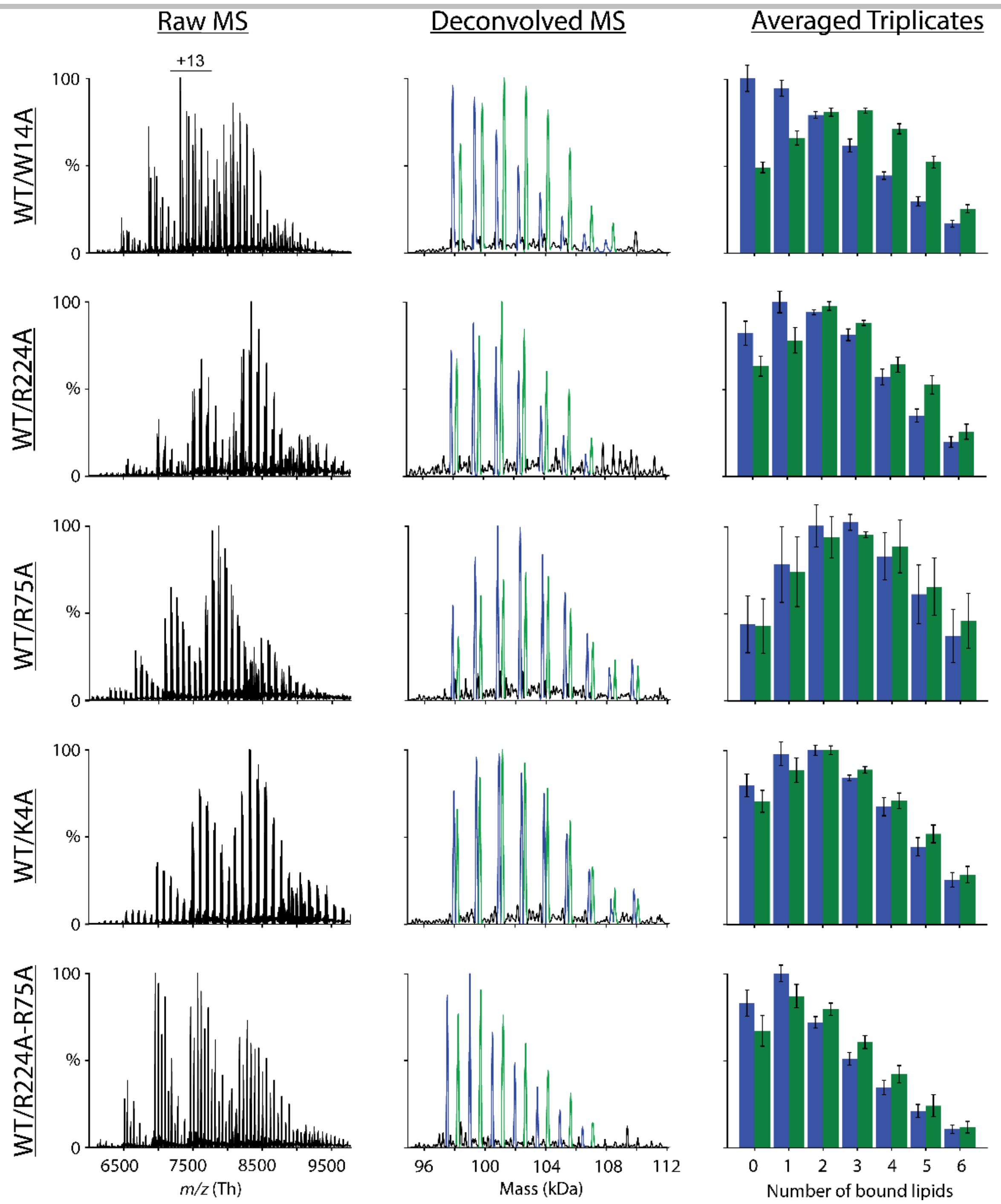

**Figure S2.** Raw (*left column*) and deconvolved (*center column*) mass spectra for a representative replicate at 25 °C for W14A (*top row*), R224A (*top middle row*), R75A (*middle row*), K4A (*bottom middle row*), and R224A_R75A (*bottom row*) mutants with wild-type protein up to six lipids bound from top to bottom, respectively. Mutant peaks are colored in *blue* whereas wild-type protein is colored in *green* in deconvolved mass spectra and in bar charts (*right column*) showing the average and standard deviation of extracted peak areas from triplicate measurements for the respective combinations.

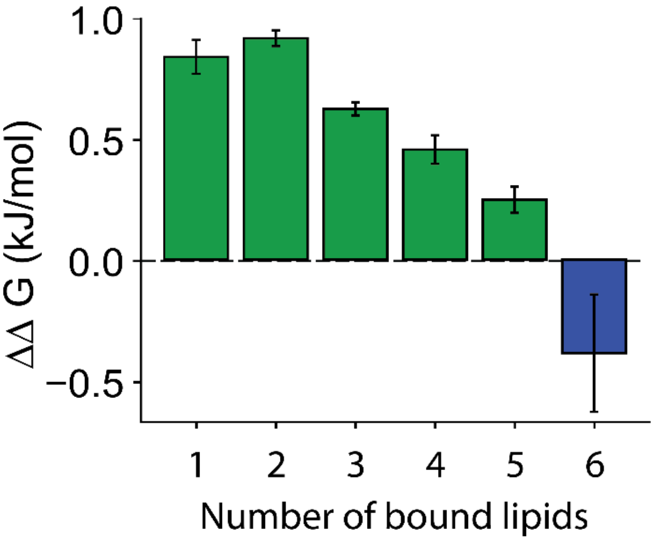

**Figure S3.** The mean difference in the Gibbs free energy change ($\Delta\Delta G$) calculated using averaged triplicate data for WT/W14A in Figure S1 for all the 6 lipids bound at 25 °C. Error bars show the 95% confidence interval. *Green* color bars represent unfavorable mutations, where CL binding is more favorable to the wild type. Negative *blue* color bars depict more favorable CL binding to the mutant.

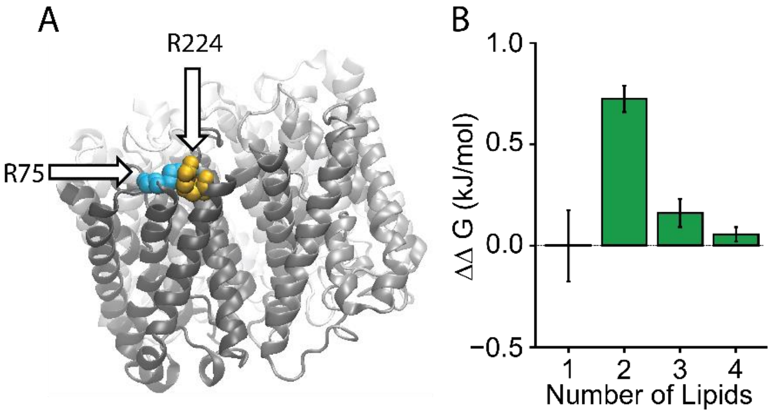

**Figure S4.** (A) AqpZ with mutant sites R224 and R75 labelled, indicated with an arrow, and colored in *yellow* and *cyan*, respectively. (B) $\Delta\Delta G$ for CL binding up to four lipids at 25 °C for R224A_R75A double mutant compared against the wild-type protein. Error bars show the 95% confidence interval.

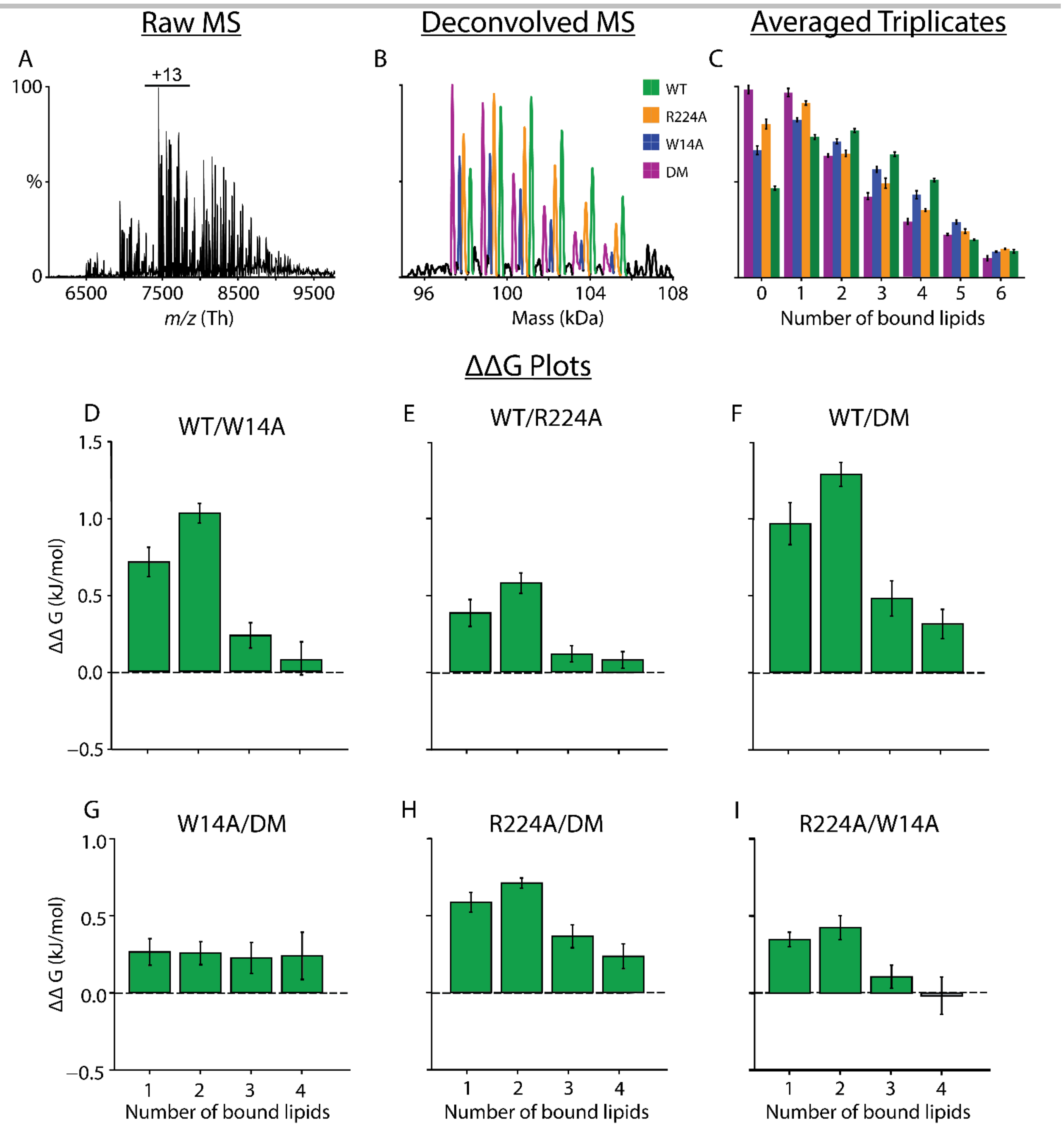

**Figure S5.** (A) Raw and (B) deconvolved mass spectra for one representative replicate at 25 °C for double mutant cycle experiment where wild type, R224A, W14A, and R224A_W14A (DM) are colored in *green*, *yellow*, *blue*, and *purple*, respectively. (C) The average and standard deviation of extracted peak area from triplicate mesaurements for each of the four species with up to six lipids bound, colored as described above. (D–I) Mean difference in the Gibbs free energy change (ΔΔG (kJ/mol)) calculated using averaged triplicate data for (D) WT/W14A, (E) WT/R224A, (F) WT/DM, (G) W14A/DM, (H) R224A/DM, and (I) R224A/W14A for 4 lipids bound at 25 °C. Error bars for D–I show the 95% confidence interval. *Green* color bars indicate that binding of CL is more favorable ot the first species in the pair. *Grey* bars represent statistically insignificant differences between binding to each species in the pair.

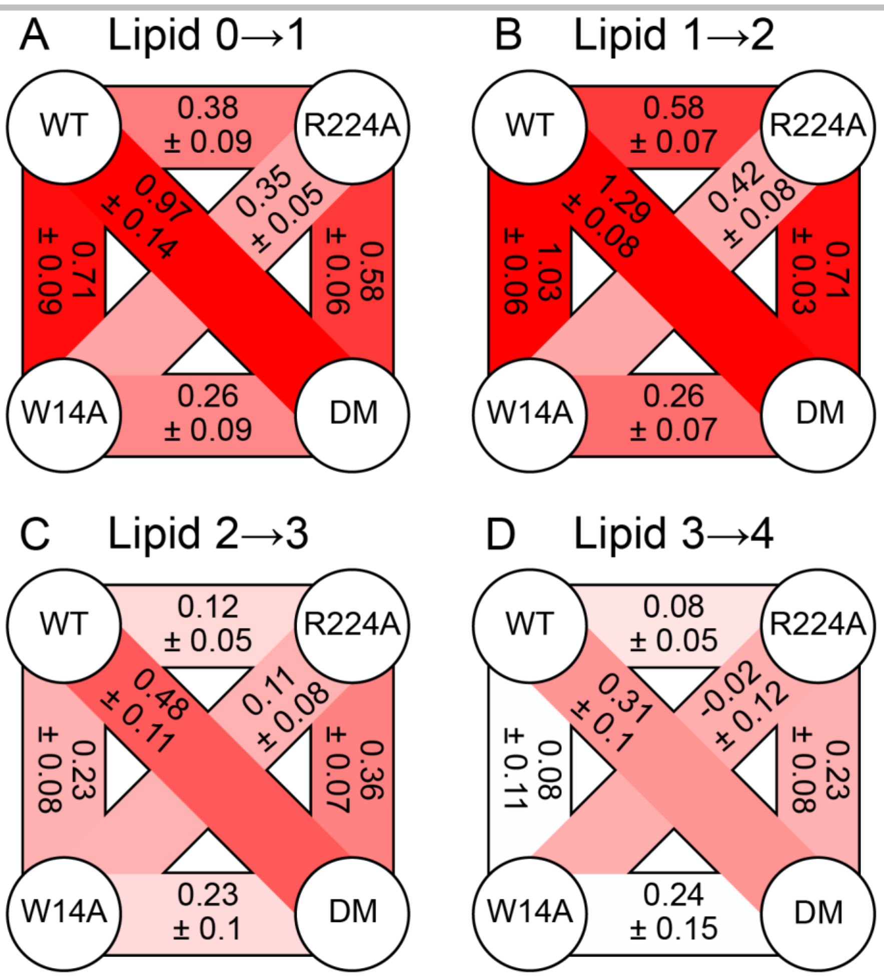

**Figure S6.** Double mutant cycles constructed with AqpZ WT, R224A, W14A, and R224A_W14A (DM) for the first four CL bindings (A–D) at 25 °C at a 1:1:1:1:200 ratio. Values indicate the average $\Delta\Delta G$ in kJ/mol from three replicate measurements with uncertainties reported as the 95% confidence interval. Statistically significant data are represented in color where positive $\Delta\Delta G$ values are depicted in *red* and negative values are indicated in *blue*. Statistically insignificant arms are in *white*. Different shades of color indicate higher and lower significance.

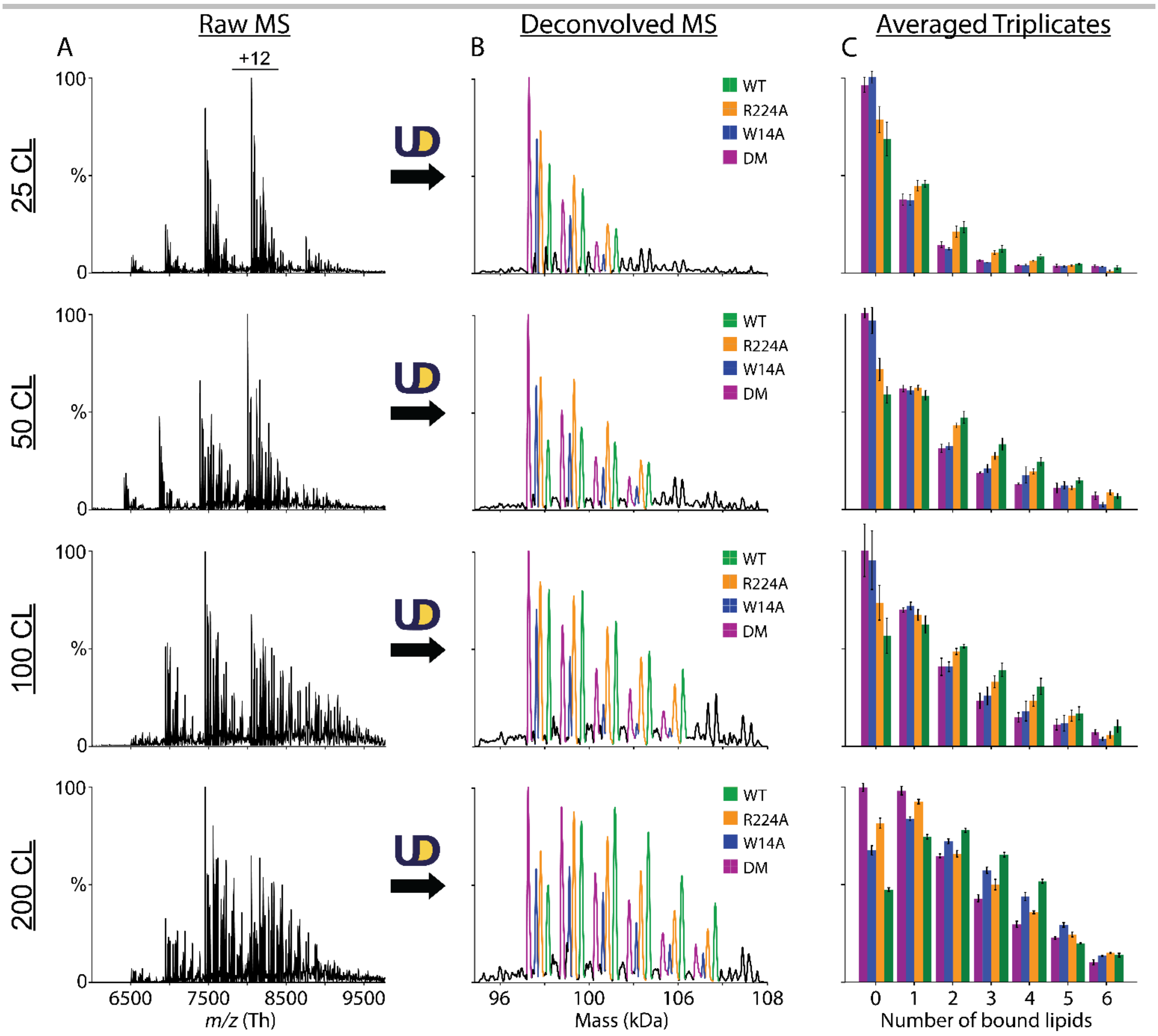

**Figure S7.** Comparison of (A) raw and (B) deconvolved mass spectra for one representative replicate at 25 °C for double mutant cycle experiment with 1:1:1:1:*n* molar ratio of WT:R224A:W14A:R224A_W14A (DM):CL, where *n* is 25, 50, 100, and 200 of CL from *top* top *bottom*. Wild type, R224A, W14A, and DM are colored in *green*, *yellow*, *blue*, and *purple*, respectively. (C) The average and standard deviation of extracted peak area from triplicate mesaurements at different CL ratios for each of the four species, colored similarly to B.

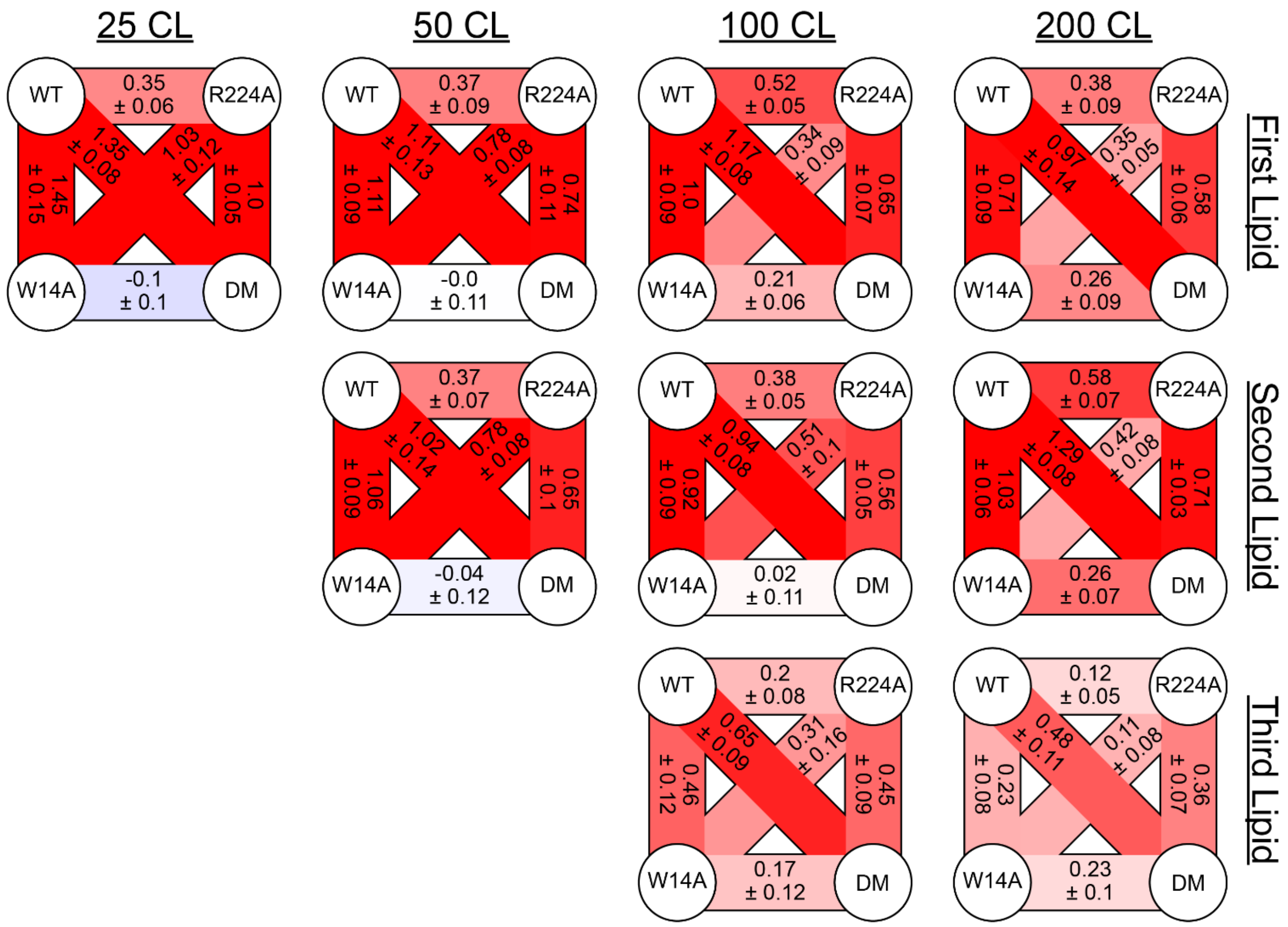

**Figure S8.** Concentration dependent experiment of double mutant cycles with AqpZ WT, R224A, W14A, and R224A_W14A (DM) mixed with CL at 1:1:1:1:*n* molar ratio, where *n* is 25, 50, 100, and 200 of CL. Double mutant cycles for up to four lipids at 25 °C at different CL ratios are shown, where values indicate the average $\Delta\Delta G$ in kJ/mol from three replicate measurements with uncertainties reported as the 95% confidence interval. Statistically non-zero data are represented in color where positive $\Delta\Delta G$ values are depicted in *red* and negative values are indicated in *blue*. Statistically insignificant arms are in *white*. Different shades of color indicate higher and lower significance. Missing panels for 25 and 50 are due to low signal for higher bound lipid states that prevents accurate measurement at higher numbers of lipids bound.

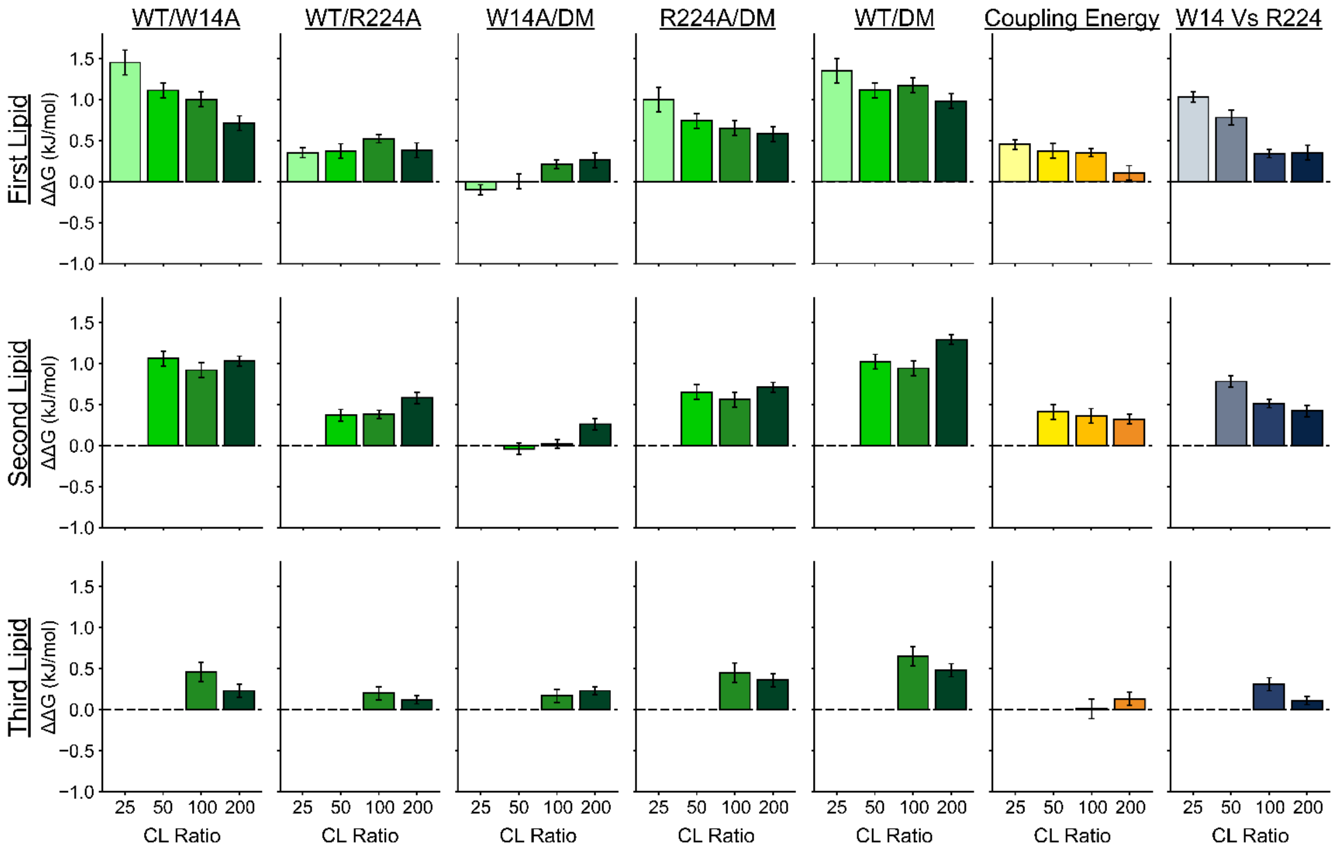

**Figure S9.** Concentration dependence of the double mutant cycles constructed with AqpZ WT, R224A, W14A, and R224A_W14A mixed with CL at a molar ratio of 1:1:1:1:*n*, where *n* is 25, 50, 100, and 200 of CL. Gibbs free energy change ($\Delta\Delta G$ (kJ/mol)) and the 95% confidence interval for CL binding between each pair-wise membrane protein combination is plotted against the CL ratio up to four lipids. Bars are color-coded in varying shades of *green*, progressing from *left* to *right* to represent increasing CL ratios. Coupling interactions between W14 and R224 residues are visually depicted in shades of *yellow*, darkening with the rising CL ratio. The relative contributions of the W14 site compared to R224, is colored similarly in shades of *blue*.

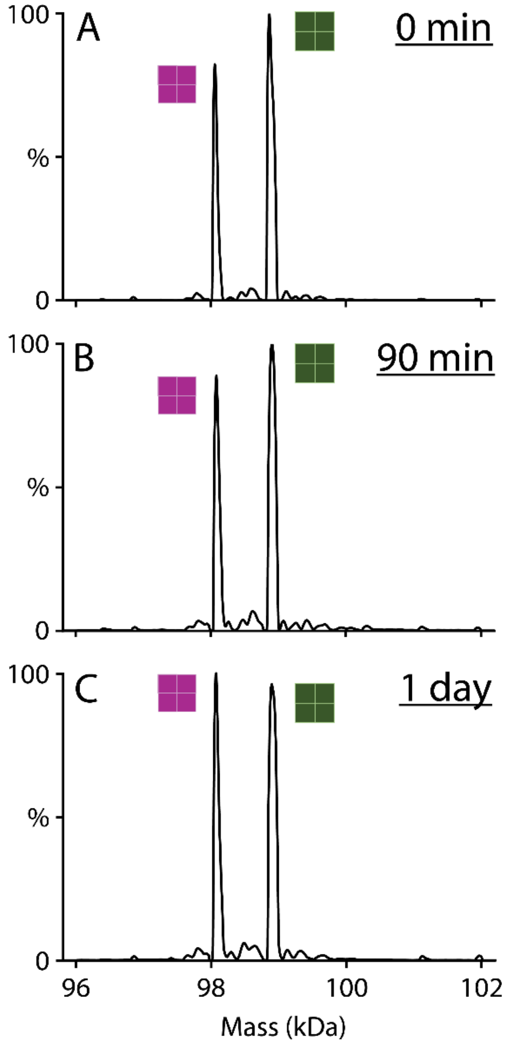

**Figure S10.** Deconvolved mass spectra of AqpZ mixed wild-type (*green squares*) and double mutant (*purple squares*) proteins without lipids incubated for (A) 0 min, (B) 90 min, and (C) 24 h. Subunit exchange between the two proteins would yield mixed mass peaks in between the two starting peaks, which are not observed. Thus, subunit exchange is negligible on the timeframe of these experiments.

### Author Contributions

H. S. Jayasekera, F. A. Mohona, and M. Ewbank performed mutagenesis, expressed, and purified proteins. H. S. Jayasekera and F. A. Mohona performed mass spectrometry (MS) experiments. H. S. Jayasekera and M. T. Marty designed the research, analyzed MS data, and wrote the manuscript.
